# Robust characterization of selectivity of individual neurons to distinct task-relevant behavioral states using calcium imaging

**DOI:** 10.1101/2024.11.30.626195

**Authors:** Huixin Huang, Garima Shah, Hita Adwanikar, Shreesh P. Mysore

**Author notes:** Co-corresponding authors with equal contribution. Co-first authors with equal contribution.

## Abstract

Investigations into the neural basis of behavior have recently employed fluorescence imaging of calcium dynamics in a variety of brain areas to measure neural responses. However, across studies, diverse and seemingly subjective methodological choices have been made in assessing the selectivity of individual neurons to task-relevant behavioral states. Here, we examine systematically the effect of different choices in the values of key parameters from data acquisition through statistical testing on the inference of the selectivity of individual neurons for task states. We do so by using as an experimental testbed, neuronal calcium dynamics imaged in the medial prefrontal cortex of freely behaving mice engaged in a classic exploration-avoidance task involving spontaneous (animal-controlled) state transitions - navigation in the elevated zero maze (EZM). We report that a number of key variables in this pipeline substantially impact the selectivity label assigned to neurons, and do so in distinct ways. By quantitatively comparing newly defined accuracy and robustness metrics for all the 128 possible combinations of levels of the key parameters, we discover in a data-driven manner, two optimal combinations that reliably characterize neuronal selectivity – one using discrete calcium events and another using continuous calcium traces. This work establishes objective and standardized parameter settings for reliable, calcium imaging-based investigations into the neural encoding of task-states.

## INTRODUCTION

Understanding the neural representations underlying distinct task-relevant states during naturalistic behavior is of major significance in systems and behavioral neuroscience. In this context, frequently asked questions are whether and to what extent individual neurons in a brain area preferentially encode one state over another. To name a few, studies have investigated whether individual neurons encode differentially social interaction vs. its absence (Frost et al., 2021; Fustiñana et al., 2021), or action-related behaviors (e.g., rear, forward, left-turn) vs. baseline status (Klaus et al., 2017), or between occupancy of safe versus aversive regions in behavioral arenas (elevated zero maze, EZM: Johnson et al., 2022; elevated plus maze, EPM: Jimenez et al., 2018; Lee et al., 2019; open field test, OFT: Grundemann et al., 2019; Jimenez et al., 2018; Johnson et al., 2022). In such investigations, the preference of each neuron for encoding one task-relevant state over another is assessed by comparing its responses to the two states in a statistically rigorous manner, often using a response selectivity ‘index’, and assigning a selectivity label to the neuron: for instance, ‘State A’-selective vs. ‘State B’- selective (Jimenez et al., 2018; Gründemann et al., 2019; Lee et al., 2019; Adhikari et al., 2011).

For such investigations, fluorescence imaging of neuronal <u>calcium</u> dynamics (“calcium imaging”) has emerged as a popular method to measure neural activity owing to its superior ability to access targeted (cell type- or projection-specific) neuronal sub-populations. However, studies employing calcium imaging to explore individual neuron selectivity have made diverse choices both in quantifying calcium dynamics and in the analytical and statistical procedures used to assess selectivity. For instance, the type of neural data employed has varied: some studies use contunous Ca2+ traces (Grundemann et al., 2019; Klaus et al., 2017; Grewe et al., 2017), others use discrete Ca2+ events (Jimenez et al., 2018), and yet others use binarized Ca2+ events (Frost et al., 2021). Whether or not neural data is binned beyond the frame rate of data acquisition has varied: no further binning (Grundemann et al., 2019; Klaus et al., 2017) versus temporally down-sampled data (Jimenez et al., 2018; Grewe et al., 2017). The shuffling methods used to assess statistical significance of the selectivity of a neuron have also varied: some studies use a random permutation approach (randperm) to generate neuronal datasets under the null hypothesis of ‘no selectivity’ (Jimenez et al., 2018) while others used a more structured, circular shifting approach (circshift) (Grundemann et al., 2019; Fustiñana et al., 2021). Diverse choices of combinations of these parameter values have the potential to influence considerably the selectivity label assigned to each neuron and to alter interpretations about the neuronal encoding properties in a brain area. Despite this critical concern, a systematic understanding of the impact of these choices is currently lacking, as is a standardized, objective approach to assess neuronal selectivity with calcium imaging. Addressing this gap is essential for promoting robust interpretations about the neural encoding of task-relevant behavioral states and for drawing reliable connections across studies.

Here, we bridged this gap by answering the two central questions: (1) What is the impact of the various parameter choices in the quantification of neural calcium activity, and in the subsequently employed analytical and statistical strategies on the assessment of neuronal selectivity for task-relevant behavioral states? (2) Which combination of parameter values is optimal, yielding reliable conclusions about neural encoding properties? To this end, we used as an experimental testbed, neural calcium dynamics imaged in the prefrontal cortex of freely behaving mice engaged in an exploration-avoidance task. This task involved navigation in a behavioral arena called the elevated zero maze (EZM) consisting of two distinct behavioral zones, and involved transitions from one to another in an entirely animal-controlled manner, as opposed to in an experimenter-controlled manner (for instance, via timed presentation of a cue or an aversive stimulus; Grewe et al., 2017; Herry et al., 2008; Laviolette et al., 2005). This was an important feature of the task because behaviors with states defined by spontaneous transitions pose additional challenges for investigating neural encoding when compared to behaviors with experimenter-controlled task states, particularly if calcium imaging is used. This, in turn, is because of the potential temporal mixing of neural calcium signals owing to the slow decay of calcium transients, which cannot be offset through task design, for instance - by extending inter stimulus intervals to allow for complete decay of the fluorescence signal, as is commonly done in experimenter-controlled tasks.

Our findings reveal that a number of key parameters including the types of neural and behavioral data used and the shuffling methods employed for statistical testing affect significantly the selectivity label assigned to individual neurons, i.e., open arm-selective, closed-arm selective, or non-selective. Moreover, based on considerations of accuracy in estimating neuronal selectivity, as well as the robustness (or consistency) in doing so, the ‘optimal’ combination of choices in methods emerged as the use of discrete Ca2+ events and head-centric behavioral data binned at the native 50 ms bins, with non-matched number of data samples from the two behavioral zones, and randperm shuffled testing. (The remaining 100+ combinations of parameters were substantially less optimal.) Separately, our results also indicated that for analyses that benefit from (or require) the use of continuous estimates of calcium dynamics as opposed to discrete events (Gründemann et al., 2019; Klaus et al., 2017; Grewe et al., 2017; Lee et al., 2019), 2s-convolved (smoothed) Ca2+ events and head-centric behavioral data binned into 1s bins, with non-matched number of data samples from the two behavioral zones and randperm shuffled testing, constituted a nearly equally optimal alternative. Taken together, results from this work establish a standardized approach that can be used generally for characterizing, using calcium dynamics, how individual neurons encode distinct behavioral states that animals spontaneously transition into and out of during natural behavior.

## RESULTS

### Study design

Our study design consisted of three steps: (a) collecting behavioral and neural data experimentally, (b) quantifying the selectivity of individual neurons to task states, and (c) systematically investigating the effect of various key parameters in the pipeline on the inferred neuronal selectivity for task states as well as identifying optimal (reliable) combination(s) of parameter choices for characterizing neuronal selectivity (Fig. 1A).

**Figure 1.**
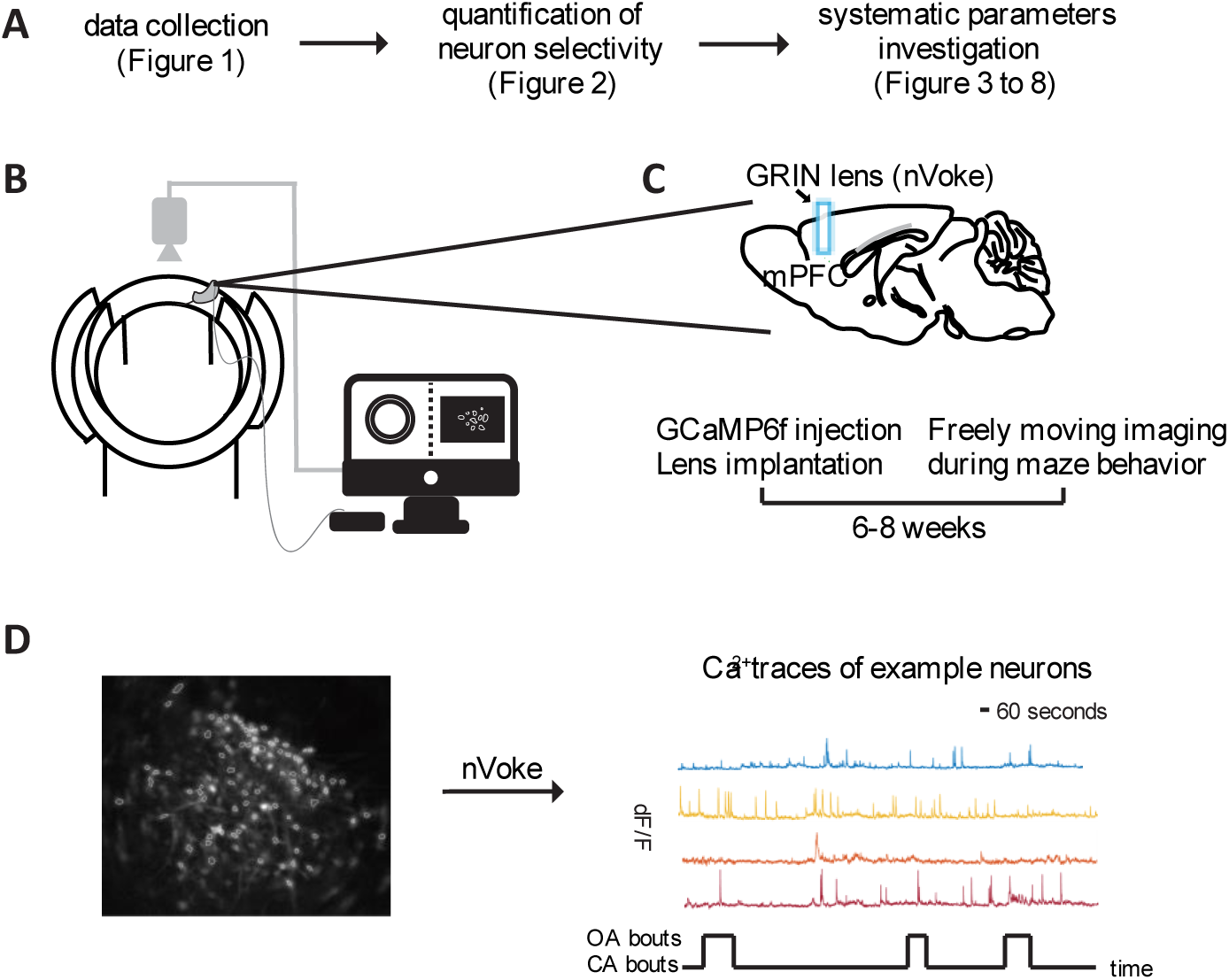
Study design. (A) Summary of the procedure for the current study. (B) (C) Schematic of the experimental setup: AAV2/5 CaMKII-GCaMP6f virus is injected into the mPFC, and a GRIN lens is implanted above the injection site. After waiting 6-8 weeks for virus expression, the animal is placed on the elevated zero maze where its behavior and the calcium activity of simultaneously imaged mPFC neurons are recorded. (D) Imaged mPFC neurons expressing GCaMP6f from an example animal (left) and calcium traces of example neurons (right) extracted from the recorded video of a 20-minute session. The time spent in open arms and closed arms is also shown.

To this end, we chose mice as our animal model, and studied spontaneous navigation by freely behaving mice in the elevated zero maze (EZM) (Shepherd et al., 1994) (Fig. 1B). The EZM consists of two distinct spatial zones – exposed, ‘open-arm’ segments representing potentially unsafe zones, and enclosed, ‘closed-arm’ segments representing potentially safe zones (Fig. 1B). Navigation in the EZM is considered to be governed by an ongoing resolution of the conflict between two innate drives: the drive to avoid unsafe spaces versus the drive to explore novel spaces. As a result, EZM navigation consists of two task-relevant states: occupancy of open arm segments, during which the exploratory drive exceeds the aversive drive, and occupancy of closed arm segments, during which the opposite is true (Shepherd et al., 1994; Montgomery, 1955). Behavioral assays involving such unconditioned navigation of potentially safe versus unsafe zones have been used extensively in the literature to study the neural basis of affective decision-making (Adhikari, 2014; Calhoon & Tye, 2015; La-Vu et al., 2020).

In mice engaged in the EZM assay, we investigated neural representations in the ventromedial prefrontal cortex (vmPFC; Fig. 1C), a brain area heavily implicated in affective decision-making (Tovote et al., 2015; Calhoon & Tye, 2015; Malezieux et al., 2023; Adhikari et al., 2011). Specifically, neural calcium dynamics in vmPFC were visualized by expressing virally the genetically-encoded fluorescent calcium indicator, GCaMP6f (Methods), and by performing cellular-resolution calcium imaging with a GRIN lens using endoscopic miniscopes (nVoke miniscope, Bruker; Fig. 1C, D) (Resendez et al., 2016). Our dataset from this study design consisted of 692 individual mPFC neurons imaged from 8 mice freely exploring the EZM.

### Characterizing the selectivity of individual neurons to task-relevant states

To characterize the selectivity of each neuron to the two task-relevant states in the EZM, namely, open arm occupancy versus closed arm occupancy, we computed a standard modulation index (Gründemann et al., 2019; Jimenez et al., 2018; Herrero et al., 2008). Termed the selectivity index, SI, it is defined as: (average neural responsiveness (Ca2+ activity) during open arm exploration – during closed arm exploration) / (average Ca2+ activity during open arm exploration + during closed arm exploration) (Methods; Fig. 2A). The values of SI range from −1 to +1, and indicate the degree to which a neuron is preferentially active during open arm occupancy (positive SI values) versus closed arm occupancy (negative SI values). Specifically, a neuron is said to be selective for one of the states if its actual SI value is significantly different from a null distribution of SI values obtained by shuffling its open arm and closed arm neural responses (Fig. 2B; Methods). The SI values and null SI distributions of three example mPFC neurons that lead us to infer, respectively, the three different types of selectivity, namely: CA-selective, non-selective and OA-selective, are illustrated in Fig. 2C.

**Figure 2.**
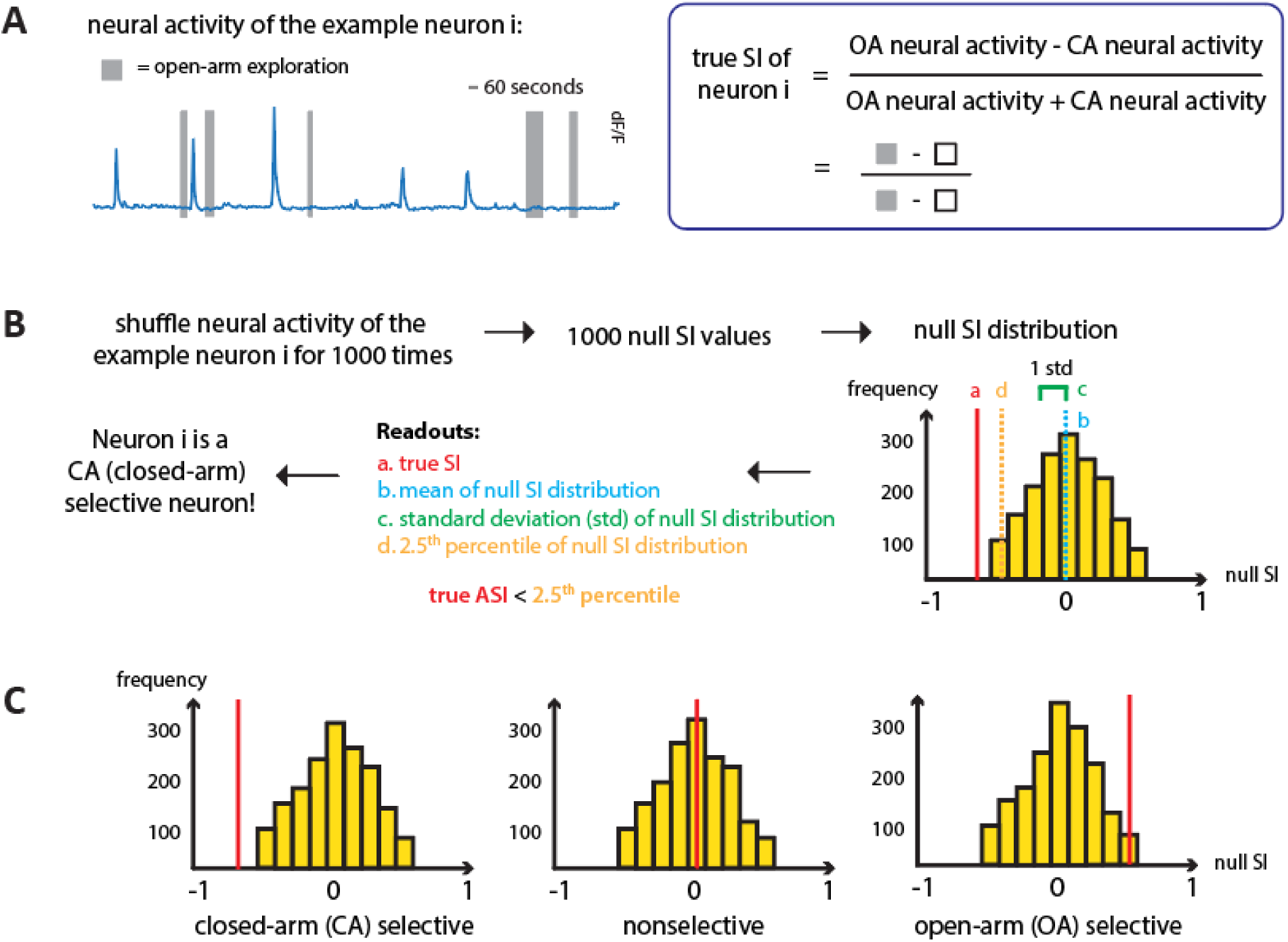
Determine selectivity of individual neurons to open-arm versus closed-arm. (A) (Left) Calcium transients of an example neuron that fired preferentially in the closed arm. The time spent in open arms is shown by gray bars. (Right) The formula for calculating the true selectivity index (SI) of an individual neuron. (B) Steps for determining the selectivity of an individual neuron. (C) SI null distribution constructed from 1000 null SI values with the true SI value of example open-arm selective, closed-arm selective, and nonselective neurons.

### Key parameters in the experimental + analysis pipeline relevant to characterizing neuronal selectivity for task-relevant states

Characterizing the selectivity of individual neurons to task states per the standard approach above necessitates apriori choices of various parameters along the data pipeline. This pipeline consists of three major stages (Fig. 3A). Here, we identify the key parameters in each stage, discuss which ones merit inclusion in our analysis, and why.

**Figure 3.**
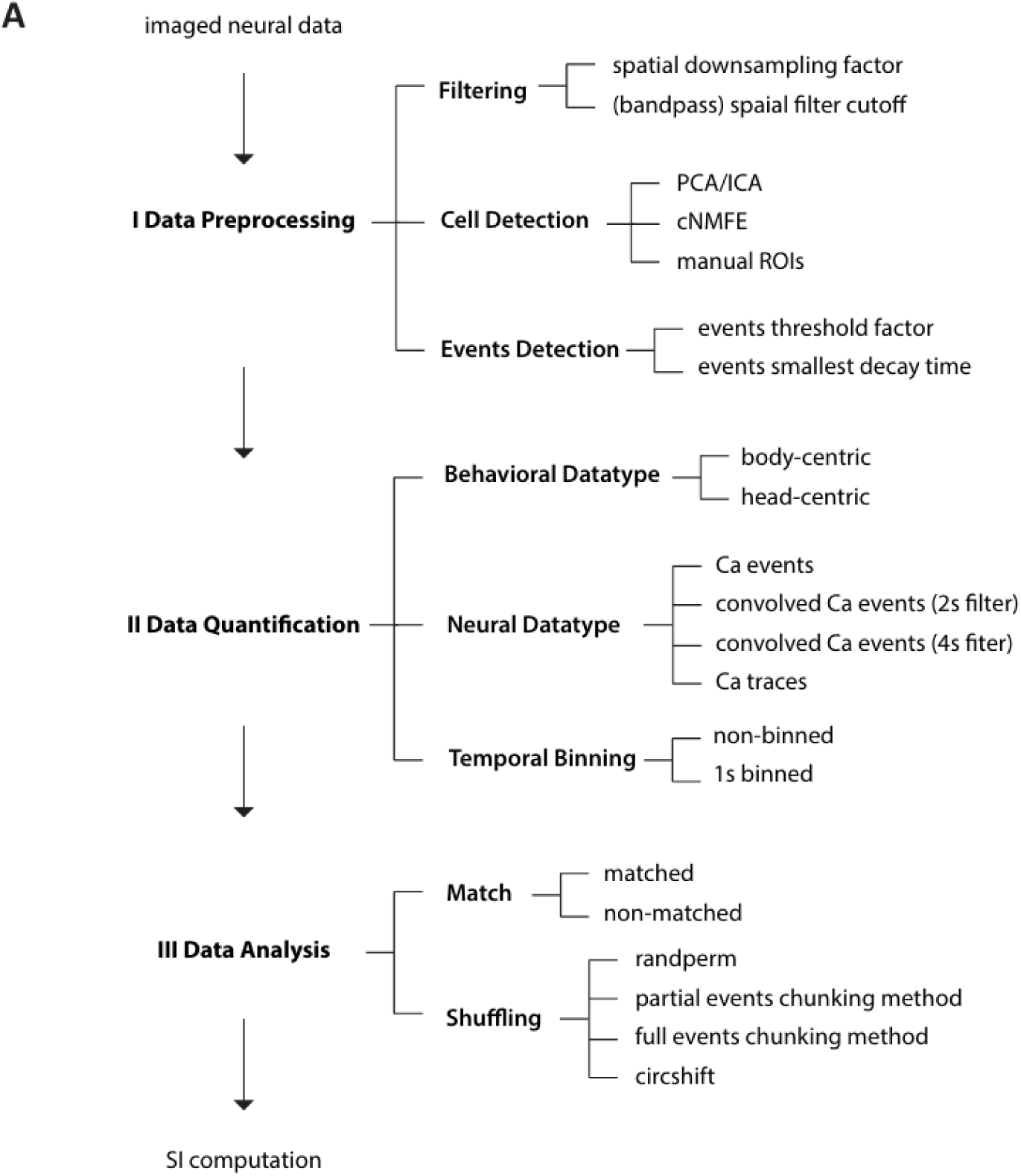
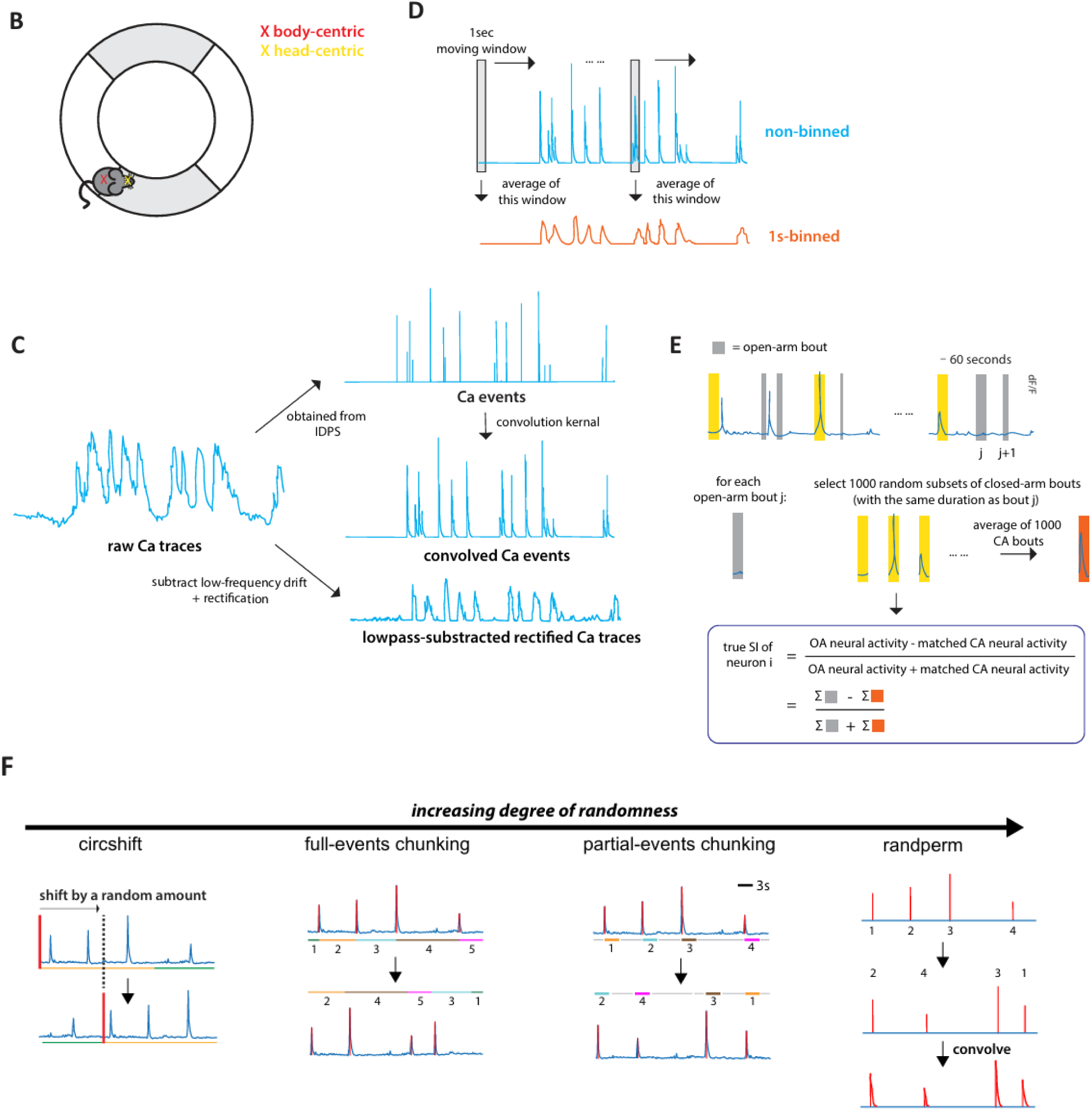
Description of the key parameters under investigation. (A) Key parameters involved in each of the three major data processing stages from data collection to assessment of selectivity. (B) Using a single coordinate to characterize the position of the mouse in the maze: head-centroid position versus body-centroid position. (C) Non-binned calcium transients (top) versus 1s-binned calcium transients (bottom). The 1s-binned calcium transients were obtained by averaging values in a moving 1-second window. (D) Illustration of the types of neural data investigated in the current study. Raw Ca traces and Ca events were preprocessed and output directly from Inscopix Data Preprocessing Software (IDPS). Convolved Ca events were obtained by applying a convolution kernel to the Ca events. Ca traces analyzed in the current study referred to the lowpass-subtracted rectified Ca traces, which were obtained by subtracting the low-pass component from the raw Ca traces and rectifying them. (E) Illustration of the number-matched method. The time spent in open arms is shown by gray bars. Yellow bars refer to random subsets of closed-arm bouts. (F) Representations of the procedures of the four shuffling methods, arranged from the lowest degree of randomness to the highest degree of randomness.

The first ‘preprocessing’ stage involves (i) spatial filtering of raw fluorescence videos obtained from the microscope, followed by (ii) identification of individual cells (with their DF/F calcium traces) from these videos, and (iii) the detection of calcium events from the DF/F trace of each cell. As described below, the parameter choices involved in all three aspects of this first stage do not significantly affect the computation of neuronal SI. For spatial smoothing, a range of standard filters are used commonly in the literature, differing in spatial downsampling factor and low/high cutoff of the filter. In theory, different filter specifications can quantitatively affect the DF/F traces of calcium activity extracted from the videos, thereby potentially affecting the eventual computation of neuronal selectivity index. In practice, however, we found that the calcium traces extracted using a range of commonly used spatial filter parameter values were highly correlated *with one another* (Fig. S1C), revealing a minimal potential impact of typical spatial filtering rules on the eventual computation of SI. For cell identification, the commonly used approaches are manual identification, PCA/ICA analysis (followed by manual verification), or cNMFE analysis (followed by manual verification). Although these different approaches often impact the number and identity of detected neurons, they do not, *per se*, impact the extracted calcium dynamics of any identified neuron, and therefore, the computation of its SI. Finally, for the detection of rapid calcium transients (or “calcium events”) from the DF/F traces of individual cells, a range of standard procedures are used commonly, differing in the event threshold factor (the threshold below which a spike is not considered a rising event) and in the event decay time factor (the minimum value of the mean lifetime of the decay of a spike). In theory, the number of events detected in a trace can depend on the rules used, thereby potentially directly impacting the computation of SI. In practice, however, we found that the time-courses of detected events were highly correlated among the commonly used procedures (Fig. S1D), revealing a minimal potential impact of typical event detection rules—on the eventual computation of SI. Consequently, the parameters involved in this first, preprocessing stage of the data pipeline were fixed to typical values for the subsequent analyses (see Methods).

The second, ‘quantification’ stage involves (i) quantifying the behavioral position of the mouse in the maze, (ii) quantifying the neural activity, and (iii) temporally binning the behavioral as well as neural data. For quantifying mouse position, the commonly used variables are ‘body’ position (centroid of the body of the mouse in each video frame) or ‘head’ position (estimated center of the head in each frame). Given that mice frequently exhibit behaviors involving head movements not captured by body position alone, such as exploratory head extensions into the open arm with the body still in the closed arm, these two variables can produce differing assessments of the task-relevant zone occupied by the mouse at any instant (and thereby, potentially affect the eventual computation of neuronal selectivity). We included both these ‘*behavioral datatypes*’ for our subsequent analysis. For quantifying neural calcium dynamics, the commonly used variables are the area-under-the-curve of continuous calcium traces, or the number of discrete calcium transients (events) detected from these raw traces. Whereas the continuous traces provide the most ‘complete’ description of calcium dynamics, because of the slow (seconds-long) decay in calcium kinetics, calcium fluorescence changes that are initiated just before the animal executes a state transition can leak into the post-transitional state resulting in potential mis-attributions (over-estimation) of area-under-the-curve activity for the second state. By contrast, discrete calcium events are much less subject to such mis-attribution errors. We included both these *‘neural activity datatypes’* for our subsequent analysis.

In addition, we introduced two others, by convolving discrete Ca2+ events with a single exponential decay kernel of time constant 2s or 4s, to mimic fast versus slow kinetics of the kind observed with GCaMP6f vs. 6s. Both these convolved (continuous) traces offer the same advantages and over discrete event traces as raw traces do for high-dimensional and state-space analyses of neural data such as greater statistical power (Cunningham & Yu, 2014), and represent less noisy approximations of the raw calcium trace. Finally, for binning (temporally down-sampling) both the calcium and behavioral data, the commonly used choices in the literature range from no binning (beyond the original sampling rate, i.e., 50 ms bins), to binning data into 1s bins. We included these two extremes (50 ms vs. 1s bins) as our two options for subsequent analyses.

The third, ‘analysis’ stage, is the one in which the SI of each neuron is computed. It involves choices with respect to two key steps: (i) balancing (or not) the number of data samples between open arm occupancy and closed arm occupancy for SI calculation, and (ii) the shuffling method employed to obtain the null distribution of SI values for statistically rigorous inference of neuronal selectivity. With respect to data balancing (or matching), because mice typically spend a greater fraction of time in the potentially safe zone (closed arm) compared to the aversive zone (open arm), an imbalance between the number of data points available for these two zones is commonplace. Such an imbalance could potentially introduce biases in the estimation of SI. To address this issue, we compared the impact on SI estimation of matching exactly the number of closed-arm data samples to the typically fewer number open-arm data samples (through random sub-sampling of closed arm data), versus not matching the number of data samples for characterizing a neuron’s selectivity (Methods; Fig. 3E). With respect to data shuffling for null distribution generation, we explored four different methods (Fig. 3F). “Randperm” and “circshift” are the most commonly used shuffling methods in the literature, referring, respectively, to randomly re-arranging neural data samples in time, versus to shifting neural time-course with respect to the behavioral time-course by a random duration. The former method fully randomizes the association between neural and behavioral data, whereas the latter injects a smaller degree of randomness to this association while retaining local (temporal) structure in the neural data. To better explore the tradeoff between fully randomizing the data vs. retaining local temporal structure, we introduced two additional shuffling methods that we refer to as “partial events-chunking” (partial EC) and “full events-chunking” (full EC) methods. In the partial EC method, we defined an event chunk as the neural signal from one calcium event to x seconds after it (x=2 for 2s convolved Ca events, x=4 for 4s convolved Ca events, x=3 for Ca events and Ca traces) to account for the decay in calcium transients, and then shuffled these chunks in time. In the full EC method, we extended this by defining an event chunk as the neural signal from one event until the next, and then shuffled these chunks in time. We examined the effect of both data balancing methods and all four of the shuffling methods on subsequent SI characterization.

Taken together, the three key parameters from the data quantification stage and two key parameters of the data analysis stage described above resulted in a total of 128 unique combinations of parameter levels for characterizing the selectivity of individual neurons (128 = 2-behavioral datatypes X 4-neural activity datatypes X 2-binning widths X 2-matching levels X 4-shuffling methods.) Henceforth, each unique combination of levels of the five parameters will be referred to as a parameter ‘setting’, with there being a total of 128 parameter settings.

### Do parameter choices matter, and if so, how?

We investigated whether the diverse combinations of parameter levels had an impact on the encoding of task-relevant states by individual neurons. Specifically, we quantified the effect of parameter settings on the selectivity label assigned to each neuron, namely OA selective, CA selective, or non-selective: an observed change in the assigned selectivity label owing simply to a different combination of parameter levels would indicate a major error in inference regarding what a neuron encodes. To this end, we computed the ‘label consistency’ between each pair of parameter settings as the percentage of neurons assigned the same labels by both settings. This produced a 128×128 matrix of (pairwise) label consistency values (Fig. 4A); we only show the upper diagonal part of the matrix because it is symmetric. Bootstrap confidence intervals were also computed for each entry.

**Figure 4.**
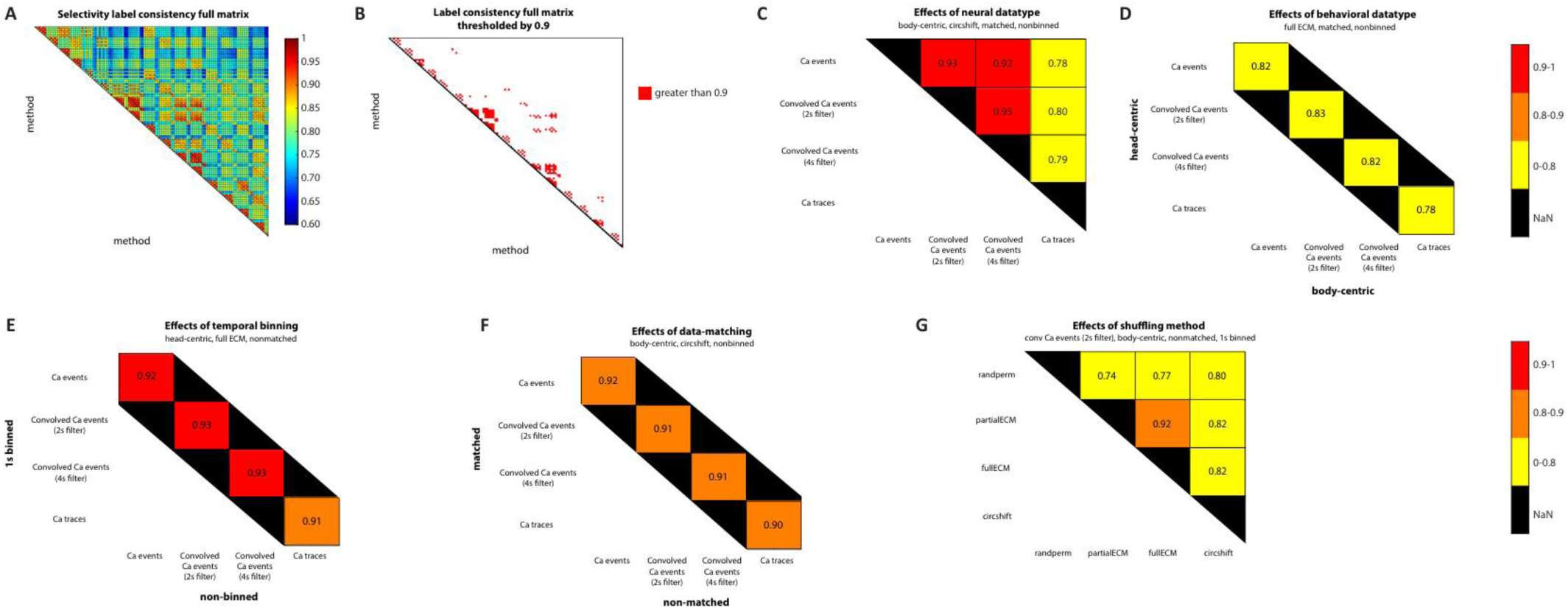
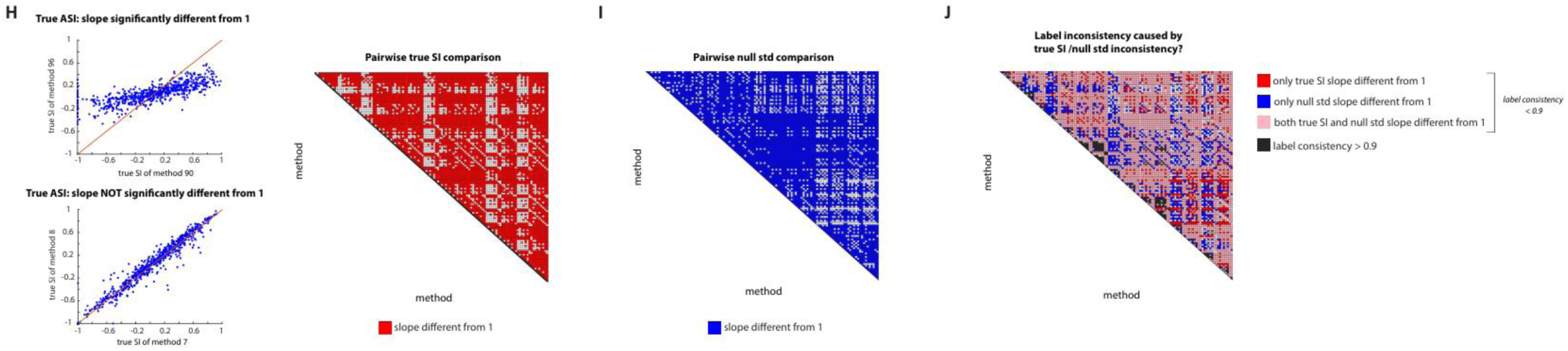
Effects of parameters on label consistency, true selectivity index (SI), and null standard deviation (STD). (A) Pairwise label consistency matrix. A 128×128 matrix visualizing the effect of parameter variations on neuron selectivity labels (OA-selective, CA-selective, or nonselective). Label consistency was defined as the percentage of neurons assigned the same label by two parameter combinations. Note that randperm does not apply lawfully to raw traces, so 8 combinations were excluded. (B) Thresholded label consistency matrix. The pairwise label consistency matrix (A) thresholded at 0.9. Bootstrap confidence intervals (CIs) were computed for each matrix entry, and the lower bound of the CI was compared to the threshold. (C-G) Submatrices of parameter effects. Example pairwise label consistency submatrices illustrating the main effects of each parameter. Colors indicate consistency levels based on CI lower bounds: red (high, >0.9), orange (medium, >0.8), and yellow (low, ≤0.8). (H) True SI consistency. Right: A 128×128 matrix of true SI consistency, computed by fitting a line to true SI values for each pair of parameter combinations. Bootstrap CIs for the slope determined whether it significantly deviated from 1. Left: Two example best-fit lines, representing low (top) and high (bottom) true SI consistency. (I) Null STD consistency. A 128×128 matrix similar to panel H, assessing null STD consistency. (J) Combined consistency thresholding. A 128×128 matrix showing true SI and null STD consistency, filtered by the thresholded label consistency matrix (panel B). Black indicates parameter pairs with label consistency >0.9. Among pairs with low label consistency (<0.9), red indicates low true SI consistency, blue indicates low null STD consistency, and pink indicates low consistency for both metrics.

Examining this matrix revealed, remarkably, that pairwise consistency of assigned labels was low for the majority of pairs of parameter settings. Nearly all (96.8%) of the entries of the label consistency matrix were lower than a nominal cut-off value of 0.9 consistency (Fig. 4B – red values). Notably, nearly three-fourths (74.3%) of the entries were ‘very low’ - lower than an even more permissive cut-off value of 0.8 consistency (Fig. S2A).

To gain a better understanding for which parameters (and their specific levels) contributed to the changes in selectivity label, we examined sub-matrices of the consistency matrix systematically such that we could assess the effect of varying the values of one parameter while keeping those of the others fixed. First, we examined the effect of varying *neural activity datatype*, a parameter from the ‘data quantification’ stage. As a starting point, when we fixed the other four parameters to the following levels: behavioral datatype = ‘body-centric’, binning width = ‘50-ms’, shuffling method = ‘circshift’, and OA-CA data matching = ‘yes’, the corresponding submatrix of consistency values revealed that label consistency was high between the levels ‘Ca events’ and ‘Ca convolved events’ (with either a 2s/4s filter), easily exceeding the nominal cut-off of 0.9 consistency (Fig. 4C-red shading). However, label consistency was very low between the level of ‘Ca traces’ and any of the other three neural activity datatype levels, falling below the lower cut-off of 0.8 consistency (Fig. 4C-yellow shading). This pattern of inconsistencies in the last column of the sub-matrix was found for most other combinations of levels of the remaining four parameters (Fig. S2B1, B2), exposing the role of *neural activity datatype*, and specifically of the level ‘Ca events’ in producing inconsistencies in inferred selectivity labels, with additional variations observed for some combinations - indicating also the presence of interactions among *neural activity datatype* and other parameters (Fig. S2B2). Thus, *neural activity datatype* parameter, and specifically, its level ‘Ca traces’, exerted a large impact on the inferred selectivity labels across combinations of levels of the other four parameters.

Next, we examined the effect of varying each of the other two parameters from the ‘data quantification’ stage. We found that *behavioral datatype* exerted a large impact on the inferred selectivity label across combinations of levels of the remaining four parameters (Fig. 4D; Fig. S2C1,C2). *Binning-width* also had an impact, though less severe, on the inferred selectivity labels (Fig. 4E, Fig. S2D1,D2).

Finally, we examined the effect of varying each of the two parameters from the ‘data analysis’ stage. We found that the *OA-CA data matching* parameter (indicating whether or not the number of open-arm and closed-arm datapoints were balanced during characterization of a neuron’s selectivity label) had an impact on the inferred selectivity labels (Fig. 4F, S2E1,E2). Additionally, the *shuffling method* parameter exerted a large impact on the inferred selectivity labels of neurons across combinations of levels of the other four parameters (Fig. 4G, S2F1,F2).

How might these parameters affect the inferred selectivity labels? The assignment of the selectivity label for a neuron depends both on the true SI value computed, and an assessment of whether it is significantly different from the null distribution of SI values. We wondered if label inconsistencies between parameter settings were attributable to differences in the true SI, in the standard deviation of the null distribution of SI, or both. To this end, we compared the true SI values computed for all neurons using one parameter setting against those computed using another setting (examples in Fig. 4H left). If they differed significantly between the two parameter settings (slope of best-fit line significantly different from 1; Methods), we concluded that a change in true SI value was a major contributing factor to label inconsistency between these two settings. Across all pairs of parameter settings, this analysis yielded another 128×128 matrix, but this time of consistency between true SI values (Fig. 4H-right). Similarly, we generated a third 128×128 matrix of consistency showing consistency between standard-deviation (STD) values of the null SI distributions. (Fig. 4I).

We then combined these two matrices, using the thresholded label consistency matrix (Fig. 4B) as a mask, to assess the contributions of true SI values and null STD values in producing inconsistencies in selectivity labels (Fig. 4J). Among the pairs of parameter settings with label consistency *lower* than the nominal 0.9 cut-off (i.e., white entries in Fig. 4B), we found that the majority showed significant changes in both true SI and null STD (Fig. 4J – pink cells), while some showed significant changes only in the true SI value (Fig. 4J – red), or only in null STD (Fig. 4J – blue). (There was a small fraction of entries that did show label inconsistency at the 0.9 cut-off level, but without significant changes in either true SI or null STD values (Fig. 4J-white): for these entries, the true SI and/or null STD did differ between the two settings, but are not significantly different under the somewhat arbitrarily chosen confidence interval levels.) Separately, among the pairs of parameter settings with label consistency *greater* than the nominal 0.9 cut-off (i.e., red entries in Fig. 4B), we found that some pairs did exhibit significant differences in the computed true SI values (and/or null STD values; Fig. 4J-black), potentially affecting other inferences about cellular encoding preferences that depend on the magnitude of SI rather than just its category.

Thus, these results revealed not only that the inferred selectivity labels of neurons are sensitive to key parameters (and interactions among them), but also that the effects of parameter levels on selectivity labels can occur through different ‘mechanisms’ (by affecting the true SI value, or the null STD, or both). Taken together, they establish the substantial, widespread, and diverse impact of parameter choices in the pipeline from calcium imaging data acquisition through analysis on the characterization of the encoding properties of individual neurons for task-relevant states.

### What is the ‘optimal’ parameter combination for analysis of neuronal selectivity?

Considering the extent of inconsistency produced by changing the levels of nearly all the key parameters, an important question that remained was: what choices of parameter levels should one use in order to make reliable inferences about the encoding properties of individual neurons? To answer this question, we adopted a standard approach from model selection literature, namely, of assessing the accuracy as well as precision (or robustness) of each model (Hastie et al., 2009), with the model exhibiting the highest accuracy and robustness taken to be the ‘optimal’ one. In our case, this would mean that the parameter setting with the highest accuracy and robustness would yield the most reliable inferences about neural encoding properties.

First, we defined the accuracy of a parameter setting as its ability to assign selectivity labels to neurons ***correctly***. Since the ground truth about neuronal labels is typically not known apriori, we developed a novel approach. We identified by inspection, so-called ‘template’ neurons as reference points for evaluation of accuracy. Template neurons were defined as those exhibiting neural activity patterns during occupancy of the OA and CA behavioral zones that were so distinctive that they could be used to incontrovertibly conclude their encoding preferences by simple visual inspection. Example template neurons that are, respectively, OA selective, CA selective, and non-selective, are shown in Fig. 5A (these neurons are from different animals). A total of 33 template neurons (7 OA-selective, 13 CA-selective, and 13 nonselective) were identified from the full set of 692 neurons. Using these template neurons, we defined the accuracy of each parameter setting as the percentage of template neurons that it labeled correctly. Results showed that only 16 out of 120 parameter settings achieved 100% accuracy (Fig. 5B); and we took these to be the superior parameter settings from the perspective of label accuracy.

**Figure 5.**
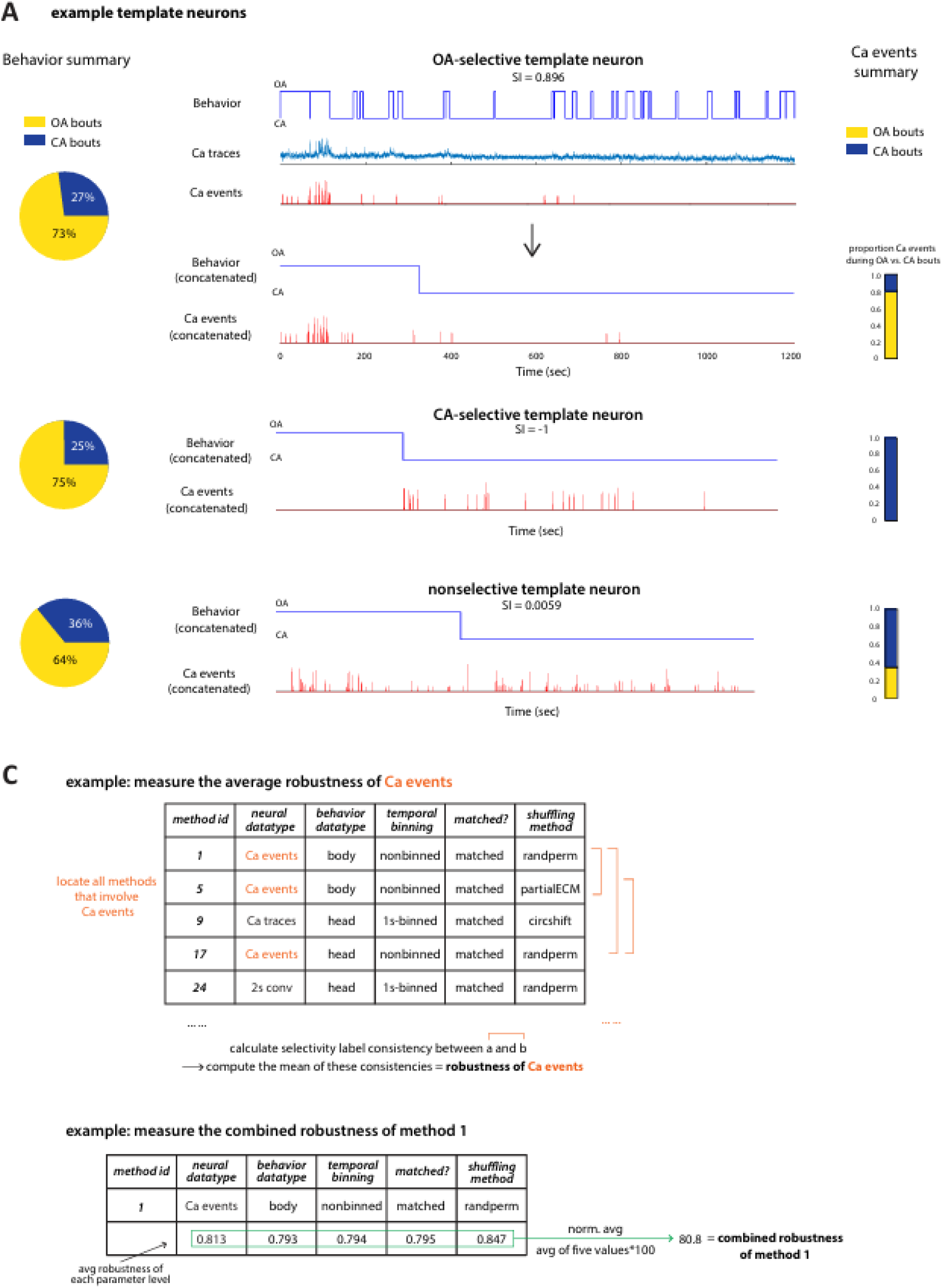
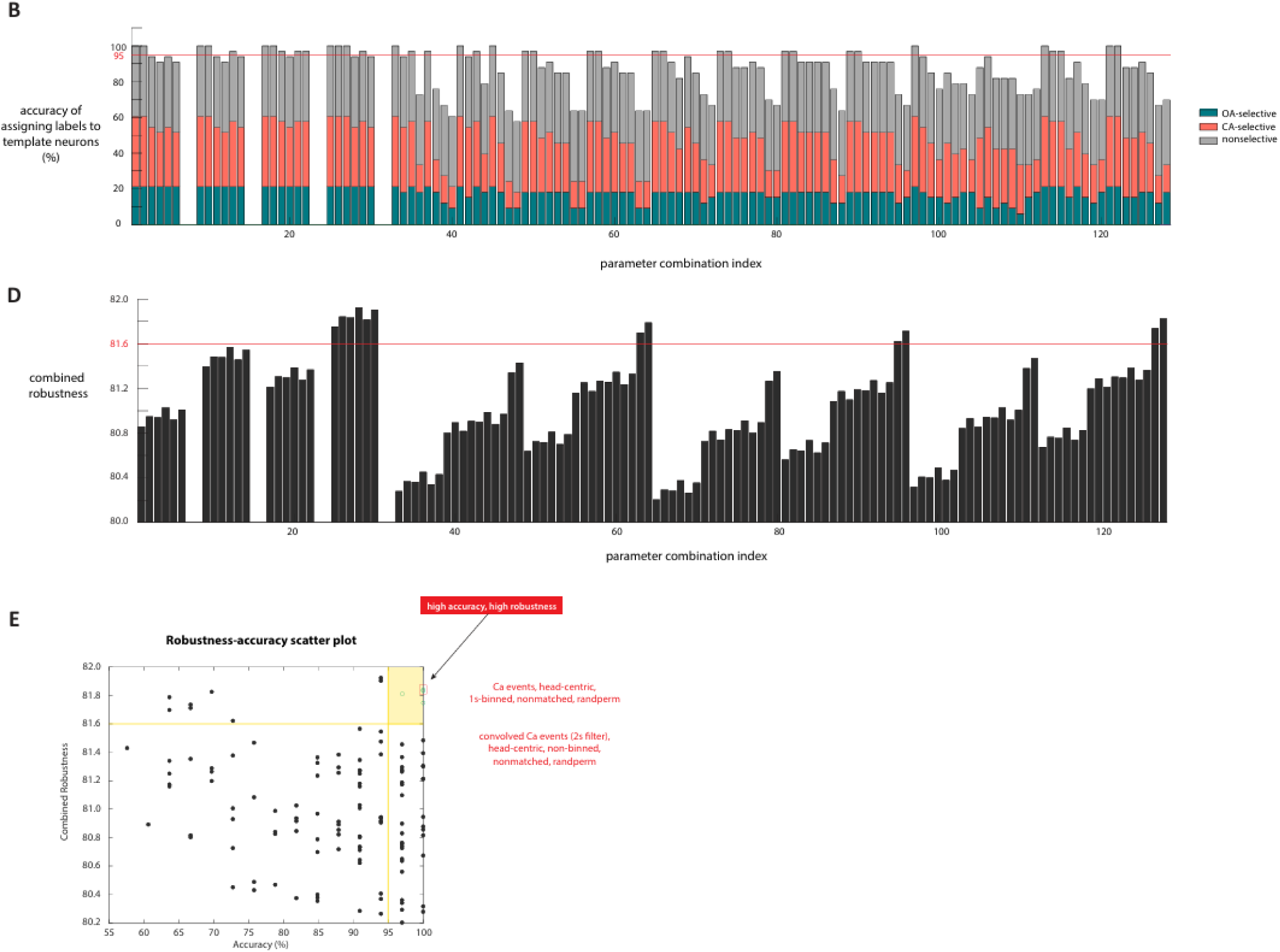
Accuracy-robustness analysis. (A) Template neuron examples. Left: Behavioral summary pie charts showing the percentage of time animals spent in OA and CA. Middle: Behavioral bouts, raw Ca traces, and Ca events for an OA-selective template neuron are shown over time. To visualize patterns across behavioral bouts, OA/CA bouts and their associated Ca events were concatenated, ensuring that events occurring during OA (or CA) bouts remain grouped. Concatenated behavior bouts and Ca events are shown for all three example template neurons. Right: Proportion of Ca events occurring during OA versus CA bouts. (B) Parameter combination accuracy. The percentage of template neurons correctly assigned to their selectivity labels for each parameter combination. (C) Robustness calculation. Top: Procedure for calculating the average robustness of a specific parameter value, using calcium events as an example. Bottom: Calculation of combined robustness across all five parameter values. (D) Combined robustness. Robustness scores for all parameter combinations. (E) Scatter plot of combined robustness versus accuracy. Yellow horizontal and vertical lines denote robustness (81.6) and accuracy (95) thresholds, respectively. Two parameter combinations with the highest accuracy and robustness were deemed optimal.

Next, we defined robustness of a parameter setting based on the extent to which each of its levels contributed to inconsistencies in the selectivity labels generated following perturbations to the levels of the *other* parameters. We reasoned that greater label inconsistency (or more variability) was less desirable, and therefore, that parameter settings with more robust constituent levels were desirable. To this end, we first defined the average robustness of each level of a parameter as the average of consistency values of all the parameter settings that contained that level (Fig. 5C, top; consistency values from Fig. 4B). This metric captured quantitatively the average extent to which the presence of that particular level of a parameter caused inconsistencies upon changes to the levels of the other parameters. Between two levels with different values of this metric, the presence in a parameter setting of the level with the lower value is more likely to cause instability or changes in the inferred selectivity, and is therefore, less desirable. (Conversely, the presence of a level of a parameter that has the highest average robustness is most desirable.)

Extending this reasoning from levels of individual parameters within a setting to parameter settings themselves, we defined the <u>combined robustness</u> of a parameter setting as the normalized combination of the robustness of each of its five constituent levels (Fig. 5C, bottom). Consequently, the parameter setting with the highest combined robustness would be the most desirable one. Our results showed that 12 out of 120 methods achieved a combined robustness greater than 81.6 (Fig. 5D).

With the accuracy and combined robustness of each parameter setting defined quantitatively, we constructed a combined robustness-accuracy scatter plot, in which each dot represents one parameter setting (Fig. 5E). The most effective settings are, by definition, the ones with high values for both accuracy and combined robustness. Four parameter settings displayed these desirable characteristics, with two standing out and designated as ‘optimal’. The top-performing combination had the following constituent levels: *neuronal activity datatype* = Ca2+ events, *behavioral datatype* = head-centric data, *temporal binning width* = 1s, *OA-CA data matching* = non-matched; *shuffling method* = randperm. This parameter setting promoted the use of discrete calcium data (with Ca2+ events). Importantly, a parameter setting involving the use of continuous calcium data also emerged as an extremely close second, with nearly identical accuracy and robustness to the optimal one. Its constituent levels were: *neuronal activity datatype* = traces of convolved Ca2+ events with 2s kernel, *behavioral datatype* = head-centric data, *temporal binning width* = 50 ms, *OA-CA data matching* = non-matched; *shuffling method* = randperm.

Thus, based on our analysis, we identify two ‘optimal’ parameter settings for inferring the encoding properties of individual neurons for task-relevant states using calcium imaging: one for discrete Ca2+ data (Ca2+ events), and another with continuous traces of Ca2+ data (2s-convolved Ca2+ events).

## DISCUSSION

We investigated the impact of choices regarding the values of key parameters in the data processing and analysis pipeline on inferences about the selectivity of individual neurons to distinct task-relevant behavioral states. We also identified the optimal parameter combinations. To achieve these goals, data obtained by imaging individual mPFC neurons in mice freely exploring the EZM served as our test bed. However, the conclusions are not unique to the specific choice of animal, or the behavioral task or brain area; they should apply generally to characterizations of individual neuron selectivity to behavioral states using calcium imaging in animals performing spontaneous state transitions.

Our findings particularly address the issues faced by studies that consist of three essential elements: characterization of individual neural code, use of calcium imaging, and the absence of experimentally controlled stimuli. Coding by neurons for different behavioral states is of major interest in systems and behavioral neuroscience. The idea of selectivity of neurons to different behavioral states is a commonly employed approach to characterize preferential coding of distinct states by individual neurons. As calcium imaging has become a powerful technique for interrogating neural activity for comparing and assessing differences in neural responses to distinct behavioral states, a key challenge with calcium imaging, compared to electrophysiological measurement of neural activity, is that calcium transients decay over relatively long periods of time (seconds). As a result, measurements of AUC to quantify responsiveness need to be mindful of the relative time courses of behavioral/stimulus events versus the time course of decay. This challenge is particularly germane in experimental designs/behavioral assays that necessarily involve purely animal-controlled behavioral transitions. In these cases, transitions can occur after a calcium event has been initiated, but before the calcium dynamics have fully decayed to baseline. Consequently, AUC calculations may lead to erroneous conclusions because a non-zero AUC during a behavioral state may actually represent residual calcium transients from an earlier behavioral state (pre-transition). By contrast, for experimental designs in which the timing of the stimuli or behavioral transitions can be experimenter-controlled stimuli (Grewe et al., 2017; Herry et al., 2008; Laviolette et al., 2005), stimulus durations and ITI can be appropriately chosen to overcome potential confounds due to the decay of calcium dynamics.

Given that discrete calcium events do not suffer from the problem of temporal leakage, it is reasonable that they should be the preferred neural data type to employ. Calcium traces, on the other hand, are suitable for high-dimensional population-level analysis (Grewe et al., 2017; Zhang & Li, 2018; Courtin et al., 2022) due to their continuous nature. However, our results indicated that the use of calcium traces produced different labels from calcium events for many neurons (Fig. 4C), potentially due to the issue of residual calcium transients. Instead, we proposed the use of convolved calcium events. The advantage of convolved calcium events is that we can control the degree of temporal leakage by varying the length of the temporal filter while retaining its continuous nature. We found that convolved calcium events with a 2s filter produced highly consistent label assignments with discrete calcium events (Fig. 4C) while other parameters were kept the same. The method which consists of convolved calcium events with a 2s filter also has the highest accuracy and precision (Fig. 5E).

We also addressed the problem of which body part to use to indicate the position of the animal in studies involving freely moving animals. When using a single coordinate to characterize the position of the mouse in the maze, the head-centroid position serves as a better descriptor than the body-centroid position. Indeed, certain behavioral gestures can only be unambiguously described using head-centric characterization. For instance, nose-pokes to peek over the edge of the open arm or head-extensions to peek across the boundary from the closed arm into the open arm are well characterized using head-center coordinates but may not be detectable using body-center coordinates as the body center may not differ between the extended versus contracted positions of the neck, i.e., between peeking versus not.

Random permutation (randperm) (Jimenez et al., 2018; Geva et al., 2023) and circular shifting (circshift) (Grundemann et al., 2019; Fustiñana et al., 2021; Frost et al., 2021) are the two most commonly used methods for obtaining the shuffled dataset. However, when applying the permutation procedure on continuous calcium transients (e.g., traces and convolved events), we need to pay particular attention to not disrupt the original structure of the signal (i.e., sharp increase followed by gradual decay when an event occurs). Circshift largely preserves the structure of the calcium signal but decreases the level of randomness. To preserve the local structure of convolved events using randperm, we need to first randperm the discrete calcium events and then apply the convolution kernel on the shuffled calcium events. However, it is difficult to find a reasonable route to apply randperm on calcium traces while retaining their original structure. To accommodate this issue, we proposed two novel shuffling methods (i.e., partial-events chunking and full-events chunking method, Fig. 3F) that can be applied to continuous calcium transients, including calcium traces. The advantages of these events chunking methods are that they can both preserve the local structure of the calcium signal while introducing more randomness than circshift.

In conclusion, our findings establish the hitherto missing foundation for objective characterization using calcium imaging of neuronal selectivity to animal-controlled behavioral states.

## METHODS

### Animals

The experiments were conducted using 8 C57BL6 mice obtained from Jackson Laboratories, Bar Harbor, Maine. The mice were allowed to acclimatize to the animal care facility for at least a week, under a 12-hour light cycle (7am to 7pm) with constant temperature (22°C) and humidity (40%), before starting any testing procedures. Food and water were provided ad libitum throughout the experiments. Experiments occurred during the standard light cycle. All protocols and animal care were in accordance with NIH guidelines for care and use of laboratory animals and approved by the Johns Hopkins University Institutional Animal Care and Use Committee.

### Surgery

We performed surgery and virus injections on mice at 10-12 weeks of age. The mouse was allowed to acclimatize to the surgery room for 30 minutes upon which it was placed in an anesthetic chamber (3% Isoflurane, 1.5 O2 level) for 5 minutes. In brief, the mouse was head fixed to a stereotactic frame (Kopf Instruments). Temperature was maintained at 36.8°C by placing a heating pad under the mouse’s body. Craniotomy was made and two injections of 350nL each of AAV1.CAMKII.GCaMP6f.WPRE.SV40 using a 0.5mL micro syringe and motorized pump (Harvard Apparatus) were lowered at a rate of 200µm/min into the mPFC region of the mouse brain (y1 = 1.86AP, x1 = 0.25ML, z1 = −2.80DV, y2 = 1.34AP, x2 = 0.25ML, z2 = −2.80DV relative to Bregma). Immediately following virus injections, a gradient index (GRIN) lens (1mmW, 4mmL, Inscopix Inc.) was implanted into the mPFC (ylens = 1.60mm, xlens = 0.25mm, zlens = 2.50mm). The microendoscopic lens was also lowered at a rate of 200µm/min into the brain and fixed in place using metabond. Upon surgical procedures, the animal was given an intraperitoneal injection of meloxicam (0.1mL), dexamethasone (0.1mL) and allowed to rest in cage under a heat lamp until recovery before returning to animal facility. Meloxicam (0.1mL) was injected once daily for the next 3 days. Mice recovered for 4-6 weeks, upon which virus expression was checked by connecting the lens to the microscope. Upon sufficient expression of the virus, the mouse was ready for mounting of the baseplate which could connect to the miniature microscope (nVista HD, Inscopix Inc.).

### Behavioral Procedures

Animals were single-housed in behaviorally enriched home cages on a 12- hour light cycle and behavioral experiments were performed during the light period. The elevated zero maze (EZM) constituted a circular platform raised above ground level (6.1cm W, 40cm inner diameter, 72.4cm above ground, Med Associates, St Albans, VT, USA) and divided into four quadrants. Two opposite quadrants were enclosed by walls of 20.3cm (“closed arms”), while the other two arms did not have any walls (“open arms”). A camera was centered above the maze apparatus to record mouse behavior. The apparatus was placed within an area surrounded by thick black curtain, lights were dimmed. Following a 30-minute acclimation to the experimental room, animals were placed in a designated open arm of the EZM, and their freely moving behavior was recorded for 20 minutes. An automated behavioral monitoring system (Ethovision, Noldus Inc v11.5, frame rate 18.96 Hz) was used to track the animal’s motion trajectories from the recorded videos. The animal was returned to its home cage and animal facility at the end of the experiment.

### Data Collection

Neural data was gathered from cellular-resolution calcium imaging of neural ensembles (50-120 neurons) in mPFC of eight mice freely moving for 20 min on an elevated zero maze (EZM). The change in fluorescence, the indication of Ca+ activity, and the behavioral video were simultaneously recorded. The two were aligned to ensure the same start and end time. The calcium imaging data was preprocessed by the Inscopix Data Preprocessing Software (IDPS), outputting the neurons identified and the neural traces and events data.

### Calcium Imaging Data Preprocessing

Raw movie data were preprocessed using the Inscopix Data Preprocessing Software (IDPS). The movies were cropped, spatially downsampled by a factor of four, high and low band pass filtered, and motion corrected. After preprocessing, PCA/ICA analysis was computed to select for individual neuronal components (see Inscopix IDPS manual for further details). Fluorescence traces for each component were extracted by the software as the average pixel intensity within the normalized ROI projected along the filtered and motion-corrected 20-Hz raw fluorescence movie. We manually inspected the traces, and those that failed to pass quality criteria (sharp peaks with an action potential like decay across 1-4 seconds time period) were excluded. We typically retained 70-90% of the components after applying the above criteria. Of the cells that we accepted, we used the traces and events data outputs from the IDPS software for further analysis. Data was imported into Matlab for analysis using custom-written software.

To derive the convolved version of the discrete Ca events, we experimented with both a 2S and a 4S spread for the decay of Ca events to convolve the Ca events data, resulting in convolved Ca events. To eliminate the low-frequency drift underlying the Ca traces, we applied a low-pass filter to the Ca traces and subtracted the slow drift component from the original data. Subsequently, we rectified the low-pass-subtracted Ca traces by setting all negative values to zero. For all analysis of the Ca traces, we utilized the rectified low-pass-subtracted Ca traces.

### Determination of neuronal selectivity for task-relevant states

Only neurons with more than 10 Ca+ events were selected for analysis. A total of 692 neurons from 8 animals were eventually included in analysis. To determine the anxiety selectivity level of each neuron, we first calculated the anxiety selectivity index (ASI) for each of them. The open-arm (OA) activity was defined as the average Ca+ activity of the neuron during the time when the animal stayed in the open arm. We used either the matched or the non-matched method to determine the closed-arm (CA) activity. In the non-matched method, CA was the average Ca+ activity of the neuron during the time when the animal stayed in the closed arm. The matched method randomly selected and matched closed-arm bouts that had the same duration as the open-arm bouts 1000 times. Each time we calculated CA activity defined as the average Ca+ activity of the neuron during closed arm bouts, and then took the average CA activity of these 1000 values, denoted as avg(CA). ASI was calculated as: 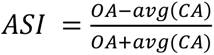.

After obtaining the ASI, a null distribution of ASI was generated for each neuron. The null distribution was obtained by shuffling the Ca+ activity for 1000 iterations. We set p=0.05 as our decision threshold, so the neurons with actual ASI greater than 97.5th of the null distribution were classified as OA selective, smaller than 2.5th of the null distribution as CA selective, and the rest as nonselective.

In addition to the raw Ca events data, we extended the application of the ASI calculation and selectivity classification procedure to the convolved Ca events data and Ca traces. The methodologies remained consistent with those described above, with the exception that the Ca events were substituted with convolved Ca events or Ca traces. Moreover, rather than employing a mere summation of discrete Ca events, we utilized the area under the curve (AUC) of convolved events to compute the average OA and CA neural activity.

### Shuffling Methods

We explored four distinct shuffling methods to obtain the ASI null distribution for characterizing the selectivity of each neuron. In the “randperm” method, we shuffled the Ca activity in time. When applying randperm to convolved Ca events, we initially shuffled the discrete Ca events and then applied the calcium-transient filter to the shuffled events to obtain the shuffled convolved events. In the “partial events-chunking” method (partial ECM), we first identified the time points at which a calcium (Ca) event occurred. We then defined each event chunk as the neural signal for x (x=2 for 2s convolved Ca events, x=4 for 4s convolved Ca events, x=3 for Ca events and Ca traces) seconds following the event, which corresponds to the approximate time needed for a Ca event to decay. These event chunks were randomly shuffled, and the time points outside of these chunks were set to zero. The “full events-chunking” method (full ECM) resembled the partial ECM, except that a chunk of event was defined as the neural signal between the occurrence of an event and the occurrence of the next event. In the “circshift” method, we shifted the Ca activity in a certain direction by a random duration.

### Calculation of label consistency across pairs of parameter combinations (‘settings’)

We evaluated 128 unique combinations of parameter values for characterizing individual neuron selectivity. These combinations were derived from five parameter dimensions: behavioral data type (2 levels), neural data type (4 levels), binning level (2 levels), matching level (2 levels), and shuffling method (4 levels) (128 combinations = 2 × 4 × 2 × 2 × 4). However, since the randperm shuffling method does not apply lawfully to raw traces, 8 combinations were excluded, resulting in 120 valid parameter combinations.

The label consistency between two given parameter combinations was defined as the percentage of neurons assigned to the same selectivity labels by both combinations. To calculate the 95% bootstrapped confidence interval (CI) for label consistency, we resampled neurons with replacement 1000 times. For each bootstrapped sample, we computed label consistency, and the 95% CI was derived from the resulting bootstrap distribution of label consistencies.

### True SI and null STD consistency analysis

To compare whether the true selectivity index (SI) or the standard deviation of the null SI distribution (null std) from two method combinations differed significantly, we used robust linear regression (*robustfit* in MATLAB) to fit a line to the true SI or null std values of neurons produced by both methods. This function uses an iteratively reweighted least-squares algorithm, which is less sensitive to outliers than ordinary least-squares regression.

We assessed whether the slope of the regression line was significantly different from 1 at a significance level of 0.01. A slope was deemed significantly different from 1 if the 99% confidence interval (CI) of its estimate did not include 1. To calculate the 99% CI, we performed 1000 bootstrap iterations, fitting a regression model to each bootstrapped sample. The 99% CI was obtained from the resulting bootstrap distribution of slope estimates. A 99% CI was used (as opposed to the 95% CI applied for thresholding label consistency) as a heuristic means to apply corrections for multiple comparisons at the 95% confidence level; this effectively provided a more conservative estimate of the number of entries with true SI or null std slopes significantly different from 1, identifying these entries as problematic.

## DATA AND CODE AVAILABILITY

Code and data used in this manuscript will be made available upon reasonable request.

## SUPPLEMENTARY INFORMATION

**Supp Figure 1.**
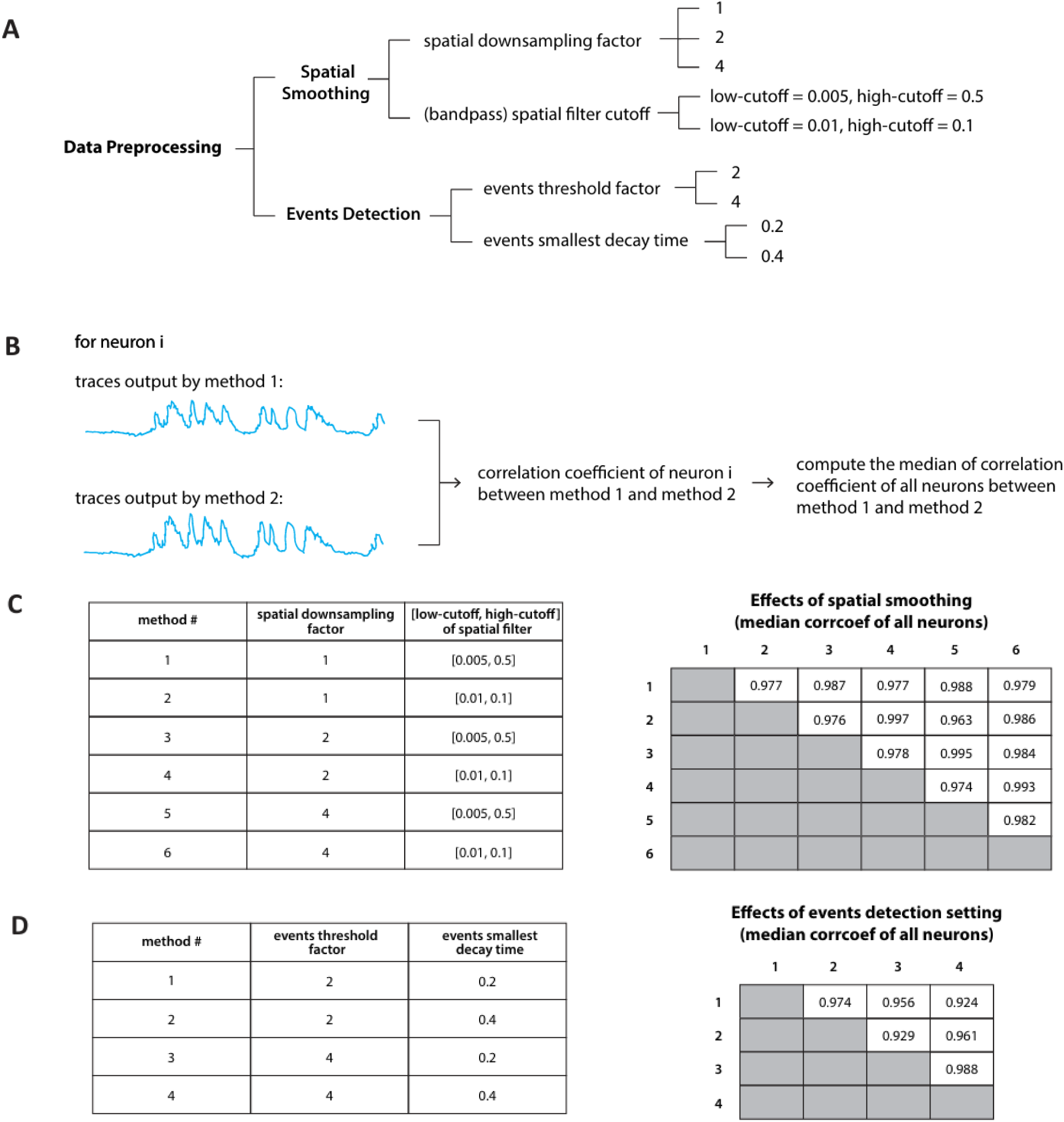
Effects of preprocessing on calcium data. (A) Key parameters in the data preprocessing stage and the standard range of choices for each parameter. (B) Procedure for inferring the overall effect of each preprocessing parameter on the output calcium data. (C) Median correlation coefficient between traces produced by methods using different parameters for spatial smoothing across all neurons. (D) Median correlation coefficient between events produced by methods using different parameters for event detection across all neurons.

**Supp Figure 2.**
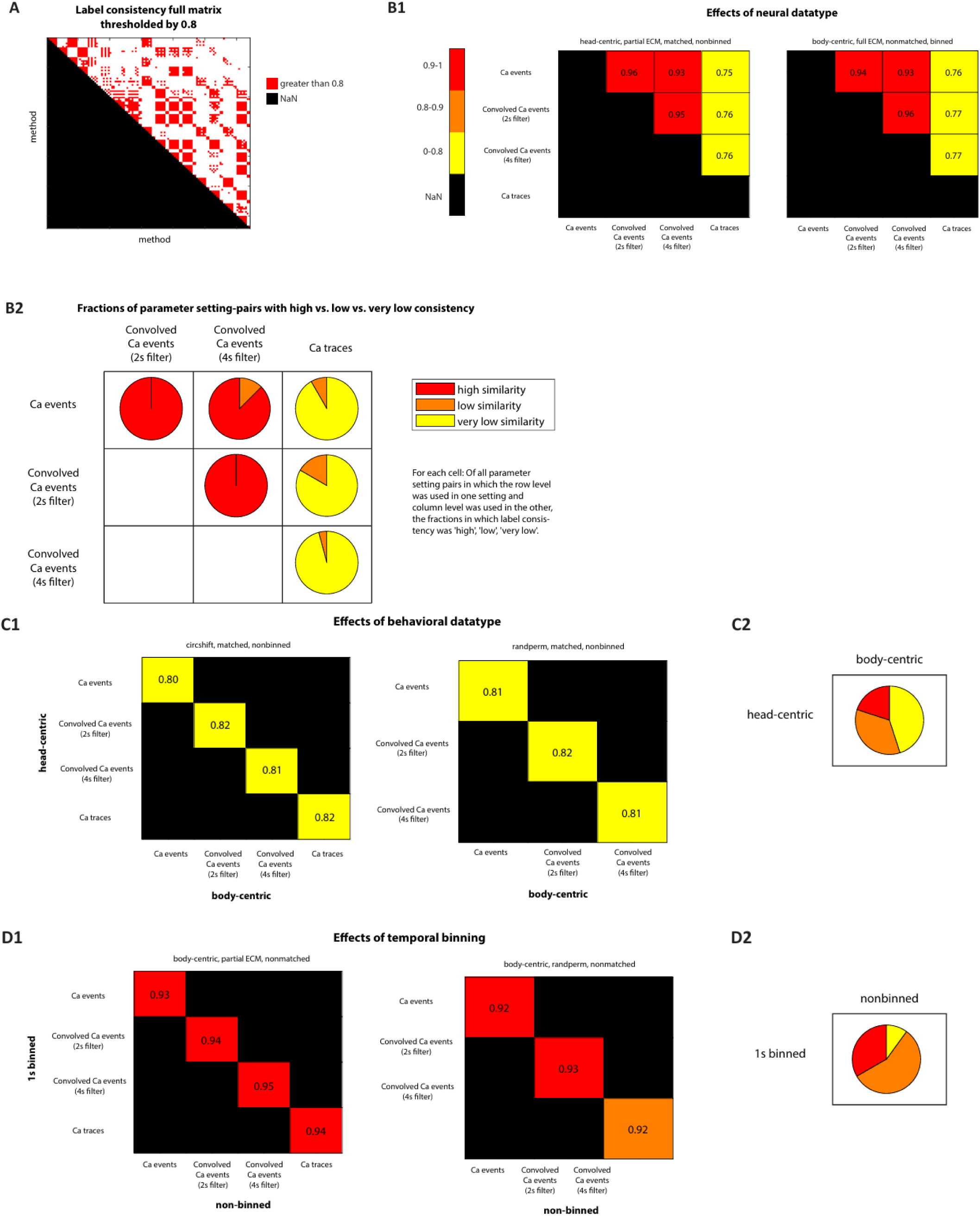
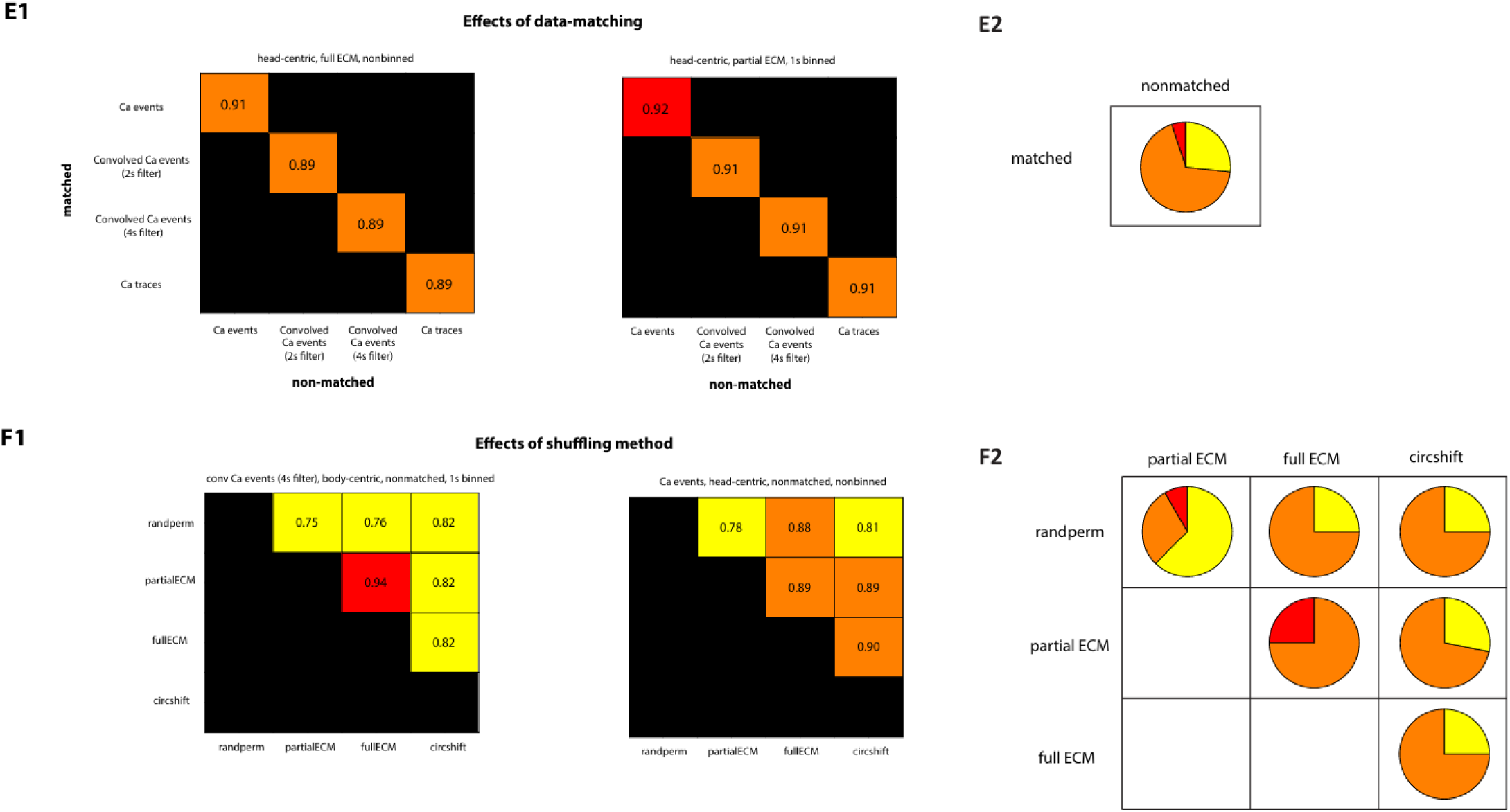
Effects of parameters on label consistency, true SI, and null std. (A) Thresholded pairwise label consistency matrix. A 128×128 matrix thresholded at 0.8 to show the effect of parameter variations on neuron selectivity labels. Bootstrap Cis were computed for each entry, and the lower bound of the CI was compared to the threshold. (B1-F1) Submatrices of parameter effects. Example pairwise label consistency submatrices illustrating the main effects of each parameter on neuron selectivity labels. Colors indicate label consistency based on the CI lower bounds: red (high, >0.9), orange (low, >0.8), yellow (very low, ≤0.8). (B2-F2) Pie charts summarizing parameter effects. For each parameter, pie charts visualize the distribution of consistency levels (red, orange, yellow) across all submatrices representing the parameter’s effect. For example, in the Ca events–Ca traces pie chart (B2), percentages of red/orange/yellow entries were calculated from 32 submatrices related to “effects of neural datatype.” The dominance of yellow and absence of red entries indicate that Ca events and Ca traces generally produce low label consistency across all conditions. Interaction effects. Pie charts also highlight interaction effects between parameters. In the absence of interactions, pie charts would display homogeneous colors (e.g., all red or orange). However, most pie charts demonstrate mixed colors, indicating significant interactions among parameters. Exceptions include the Ca events–Convolved Ca events (2s filter) pie chart and the Convolved Ca events (2s filter)– Convolved Ca events (4s filter) pie chart, which display homogeneous patterns, suggesting minimal interaction effects in these cases.

## Notes

### Competing Interest Statement

The authors have declared no competing interest.

